# IMPLICON: an ultra-deep sequencing method to uncover DNA methylation at imprinted regions

**DOI:** 10.1101/2020.03.21.000042

**Authors:** Tajda Klobučar, Elisa Kreibich, Felix Krueger, Maria Arez, Duarte Pólvora-Brandão, Ferdinand von Meyenn, Simão Teixeira da Rocha, Melanie Eckersley-Maslin

## Abstract

Genomic imprinting is an epigenetic phenomenon leading to parental allele-specific expression. Dosage of imprinted genes is crucial for normal development and its dysregulation accounts for several human disorders. This unusual expression pattern is mostly dictated by differences in DNA methylation between parental alleles at specific regulatory elements known as imprinting control regions (ICRs). Although several approaches can be used for methylation inspection, we lack an easy and cost-effective method to simultaneously measure DNA methylation at multiple imprinted regions. Here, we present IMPLICON, a new high-throughput method measuring DNA methylation levels at imprinted regions with base-pair resolution and over 1000-fold coverage. We initially designed IMPLICON to look at ICRs in adult tissues of inbred mice. Then, we validated it in hybrid mice from reciprocal crosses for which we could discriminate methylation profiles in the two parental alleles. Lastly, we developed a human version of IMPLICON and detected imprinting errors in embryonic and induced pluripotent stem cells. We also provide rules and guidelines to adapt this method for investigating the DNA methylation landscape of any set of genomic regions. In summary, IMPLICON is a rapid, cost-effective and scalable method, which could become the gold standard in both imprinting research and diagnostics.

## INTRODUCTION

Genomic imprinting describes the parent-of-origin dependent monoallelic expression of approximately 100-200 genes in mammals (reviewed in (1)). As a consequence, the inherited set of maternal and paternal chromosomes are not equivalent and they are both required for full-term development (2,3). This effect is allocated to specific chromosomal regions (4), which later were discovered to contain imprinted genes (5–8). Imprinted genes were first identified as important regulators of foetal growth and development (reviewed in (9,10)) and later shown to be involved in several postnatal endocrine and metabolic pathways, as well as in neuronal functions affecting behaviour and cognition (reviewed in (1)). Not surprisingly, genetic or epigenetic disturbances resulting in altered dosage of imprinted genes lead to severe developmental, neurological and metabolic diseases in humans [reviewed in (11,12)), such as the Prader-Willi (PWS) (OMIM#176270) and Angelman (AS) (OMIM#105830) syndromes caused by defects in the paternal or maternal chr15q11-q13 region, respectively (13,14).

Most imprinted genes are located in clusters throughout the genome, containing a *cis*-acting CpG-rich DNA element referred to as imprinting control region (ICR) (reviewed in (1)). The ICR is epigenetically marked by DNA methylation in a parent-of-origin fashion, which correlates with expression and/or silencing of the surrounding imprinted genes. Deletions of ICRs result in the loss of parental allele-specific expression within an imprinted cluster (15,16). ICRs acquire parental-specific DNA methylation in the germline, which are maintained throughout development and adulthood, resisting the global wave of demethylation and *de novo* methylation steps during early embryonic development (reviewed in (1,17,18)). The preservation of parental allele-specific methylation at ICRs, also known as germline differentially methylated regions (gDMRs), is fundamental for the correct maintenance of imprinted expression throughout life.

Despite their importance, there is currently no robust, cost-effective and high-throughput method to assess the methylation status of ICRs across multiple imprinted regions (Suppl. Table 1; reviewed in (19)). The traditional way of measuring DNA methylation is bisulfite sequencing (20). Bisulfite treatment of DNA results in deamination of unmodified cytosines to uracils, whereas methylated cytosines remain unchanged. This is followed by PCR amplification of a region of interest followed by subcloning and Sanger sequencing. This method is laborious and not cost-effective for multiple samples or viewpoints. Alternatively, bisulfite conversion can be followed by pyrosequencing analysis that provides an easier method to analyse a few (<5) CpGs within ICRs, but is not high-throughput (21). Bisulfite-treated DNA can also be converted into a next-generation sequencing library to give genome-wide information. For this approach at least 10-fold genomic coverage is required to guarantee sufficient reads at ICRs (22,23). Alternative methods for measuring methylation are reduced representation bisulfite sequencing (RRBS), that enriches for CpG-rich regions of the genome including imprinted regions (24,25), and array-based methods, as the Illumina Infinium methylation BeadChip (26). These methods, however, take several weeks and are not manageable to be routinely performed at scale. In addition, there are some bisulfite-free approaches to measure methylated DNA. The genome-wide method Methylated DNA Immunoprecipitation sequencing (MeDIP-seq) (27,28) has a low base-resolution at high costs and long preparation time. More recently, long read nanopore sequencing has successfully detected direct parental allele-specific methylation on long stretches of DNA (29). However, the potential of this technology is still hindered by the limited number of reads and its long and difficult library preparation.

Here, we present IMPLICON, a novel ultra-deep sequencing method to robustly measure DNA methylation levels with base-pair resolution at imprinted regions. This method uses bisulfite-treated DNA to generate amplicon sequencing libraries covering the majority of murine and human imprinted regions. This way, IMPLICON generates base-resolution datasets with over 1000-fold coverage that can be quickly and easily analysed to determine genomic imprinting fidelity in less than 6 days. Furthermore, we provide rules for designing additional primer sequences, making this method easily adaptable to analyse DNA methylation patterns at other genomic regions of interest. We expect that this rapid, scalable and cost-effective ultra-deep sequencing method will become a powerful tool for both imprinting research and diagnostics.

## MATERIALS AND METHODS

### Biological material

Inbred mouse genomic DNA samples were obtained from the Babraham Institute C57BL/6 J/Babr Ageing Mouse Colony as previously described (30). Genomic DNA samples from F1 hybrid animals were obtained from BL6 x CAST reciprocal crosses from the iMM JLA Rodent Facility. Animals were housed in a maximum of four per cage in a temperature-controlled room (24°C) with a 12-hr light/dark cycle. Animals were fed standard CRM (P) VP diet and both food diet and water were available *ad libitum*. All experiments involving mice were carried out in accordance with the UK and Portugal Government Home Office licensing procedures.

Genomic DNA from human peripheral blood was collected from two healthy female volunteers via fingerprick. Human ESC genomic material was collected from H9-KN2 *Nanog-Klf2* ESCs (31) and cultured in 6-well dishes under naïve (N2B27 supplemented with human LIF, 1mM Chir, 1mM PD03 and 2mM Go6983 on MEF feeder cells) and primed (Vitronectin in E8 media) conditions as previously described (32). Genomic material was also obtained from human primary fibroblasts (AS Fib. and Ctrl Fib.) and respective iPSC (Ctrl D, Ctrl E, AS A, AS B, AS D and AS E) lines from an Angelman patient and sex- and age-matched healthy individual as previously described (33). Briefly, primary fibroblasts were maintained in DMEM supplemented with 10% fetal bovine serum, 1mM L-Glutamine, and 100units/ml penicillin, 100μg/ml streptomycin (Life Technologies). iPSCs were cultured in mTeSR1 medium (STEMCELL Technologies) supplemented with 50 units/ml penicillin, 50μg/ml streptomycin (Life Technologies) in Matrigel (Corning)-coated plates. All cell lines grew in a humid incubator at 37°C with 5% (vol/vol.) CO_2_.

### DNA extraction and bisulfite treatment

Genomic DNA was isolated using either conventional phenol:chloroform:isoamyl alcohol extraction, the DNeasy Blood and Tissue Kit (Qiagen) or the AllPrep DNA/RNA Micro Kit (Qiagen) according to manufacturer’s instructions and eluted into TE buffer or H_2_O. 1μg of genomic DNA was bisulfite converted using the EZ DNA methylation Gold kit (Zymo Research) according to manufacturer’s instructions with either magnetic bead or column clean-up and eluted in 66μl elution buffer to obtain a final concentration of ~15ng/μl bisulfite converted DNA.

### IMPLICON primer design and testing

Genomic coordinates for murine ICRs or other differentially methylated regions of interest (gDMRs or somatic differentially methylated regions – sDMRs) were obtained from https://atlas.genetics.kcl.ac.uk and validated using in-house DNA methylation datasets. Appropriate SNPs in the vicinity of ICRs were acquired either from the literature (34) or from https://www.sanger.ac.uk/sanger/Mouse_SnpViewer/rel-1505. Human imprinting genomic coordinates for gDMRs were defined using oocyte and sperm methylomes (35). Genomic DNA sequences of the regions of interest were obtained from UCSC Genome Browser (genome.ucsc.edu) and imported into MethPrimer (https://www.urogene.org/methprimer/) (36) or BiSearch (http://bisearch.enzim.hu/) (37). For each region, at least 2 primer pairs for bisulfite sequencing PCR were designed, selecting those with smaller product size (optimal size 300bp, max 430bp), a minimum of 5 CpGs in the PCR product, and, no CpGs within the PCR primers. The following sequence was added to the forward (CTACACGACGCTCTTCCGATCT) and reverse (TGCTGAACCGCTCTTCCGATCTNNNNNNNN) primers (where N denotes a random nucleotide to generate a unique molecular identifier). Primer pairs were tested on 2ng bisulfite-treated genomic DNA, 0.3μM forward and reverse primer and 2x KAPA HiFi Uracil+ ReadyMix with the following conditions: initial denaturation at 95°C for 5min, 30-35 cycles of 98°C denaturation for 20 seconds, variable annealing temperature for 15 seconds and extension at 72°C for 60 seconds; followed by a final extension at 72°C for 10 minutes. The annealing temperature was tested between 60°C and 72°C. PCR products were run on a 1% agarose-TAE gel and those yielding a single strong band were selected for inclusion in the amplicon assay. Approximately 50% of designed primers yield a single strong band under these conditions. The use of 2x KAPA HiFi Uracil+ ReadyMix is crucial to ensure efficient amplification despite the lower complexity of bisulfite treated DNA.

### IMPLICON library preparation

The IMPLICON protocol consists of 2 PCR reactions. In the first reaction each sample is amplified with each primer pair in individual reactions: 30ng (2μl of 66μl eluted) of bisulfite-treated DNA) is amplified with 1.2μl of a 10μM primer pool (final 1.5μM), containing both forward and reverse primers, and 4μl of 2x KAPA HiFi Uracil+ ReadyMix in a final volume of 8μl. The hybrid mouse samples were processed in a final volume of 16μl. DNA was amplified using the following conditions: initial denaturation at 95°C for 5min, 30 cycles of 98°C denaturation for 20 seconds, variable annealing temperature for 15 seconds and extension at 72°C for 60 seconds; followed by a final extension at 72°C for 10 minutes. Annealing temperatures for each primer pair were optimised as described above and are listed in Suppl. Table 2. All PCR reactions for each individual sample were pooled together and cleaned-up using 1.5x AMPure XP beads and eluted in 20μl H_2_O. In the second PCR reaction, barcoded Illumina adapters are attached to the pooled PCR samples ensuring that each sample pool receives a unique reverse barcoded adapter. The 20μl PCR pool was amplified using 1μl of 10μM Illumina PE1.0 primer (same for all samples), 1μl of 10μM Illumina iTAG primer (distinct for each sample) and 25μl 2x KAPA HiFi Uracil+ ReadyMix in a 50μl reaction using the following conditions: initial denaturation at 98°C for 45 seconds, 5 cycles of 98°C denaturation for 15 seconds, 65°C annealing for 30 seconds and extension at 72°C for 30 seconds; followed by a final extension at 72°C for 5 minutes. Reactions were cleaned-up with 1x AMPure XP beads and eluted in 20μl H_2_O. Libraries were verified by running 1:30 dilutions on an Agilent bioanalyzer. Note that the profile of these libraries is spikey due to their amplicon nature (Supp. Fig. 1C). Libraries were sequenced using the Illumina MiSeq platform to generate paired-end 250bp reads using 10% PhIX spike-in as the libraries are of low complexity.

### IMPLICON sequencing analysis

Data was processed using standard Illumina base-calling pipelines. As the first step in the processing, the first 8 bp of Read 2 were removed and written into the readID of both reads as an in-line barcode, or Unique Molecular Identifier (UMI). This UMI was then later used during the deduplication step with “deduplicate bismark --barcode mapped_file.bam”. Raw sequence reads were then trimmed to remove both poor quality calls and adapters using Trim Galore (v0.6.2 for hybrid mouse tissues, v0.4.4 for human and inbred mouse tissues, www.bioinformatics.babraham.ac.uk/projects/trim_galore/, Cutadapt version 2.3 for hybrid mouse tissues, v1.9.1 for human and inbred mouse tissues, parameters: --paired). Trimmed reads were aligned to the mouse or human reference genome in paired-end mode. Alignments were carried out with Bismark v0.14.4 for hybrid mouse tissues and v0.18.2 for human and inbred mouse tissues (38). CpG methylation calls were extracted from the mapping output using the Bismark methylation extractor (v0.22.1 for hybrid mouse tissues, v0.18.2 for human and inbred mouse tissues). Deduplication was then carried out with deduplicate_bismark, using the --barcode option to take UMIs into account (see above). For hybrid mouse strain experiments, the data was aligned to a hybrid genome of BL6/CAST (the genome was prepared with the SNPsplit package (v0.3.4, https://github.com/FelixKrueger/SNPsplit). Following alignment and deduplication, reads were split allele-specifically with SNPsplit. Aligned read (.bam) files were imported into Seqmonk software (http://www.bioinformatics.babraham.ac.uk/projects/seqmonk) for all downstream analysis. Probes were made for each CpG contained within the amplicon and quantified using the DNA methylation pipeline or total read count options. Downstream analysis was performed using Excel and GraphPad.

From the raw data deposited in GEO under the accession number GSE146129, the reads mapped to the following murine (mm10) and human (hg38) genomic coordinates were excluded for consideration in this article for one of the following reasons: (1) failure to reach the coverage threshold (>100); (2) clear sequencing bias towards the methylated or unmethylated amplicons or to one of the SNPs, (3) regions out of the scope of this article. For inbred mice data: Chr1:63264732-63264796, Chr2:152686485-152686582, Chr2:174328905-174329102, Chr6:4746303-4746438, Chr6:58906821-58907146 and Chr17:12742173-12742420; for hybrid mice data: Chr6:4746303-4746438, Chr18:12973031-12973038 and Chr18:36988436-36988740; for human data: Chr19:16555181-16555319, Chr3:181712902-181713043 and Chr20:37521191-37521391.

## RESULTS

### IMPLICON design

To surpass the current limitations for methylation analysis at imprinted regions, we devised a bisulfite-treated amplicon next-generation sequencing (NGS) approach that we named IMPLICON (Fig.1A-C). We designed primers targeting well characterised murine imprinted regions (https://atlas.genetics.kcl.ac.uk; see Materials and Methods) and validated them as giving a single product on bisulfite-treated genomic DNA from mouse embryonic stem cells (ESCs) or tissue samples (Suppl. Fig.1A-B), resulting in 15 primer pairs covering 9 murine imprinted clusters (Fig. 1A; Suppl. Table 2). The following rules were used for designing the primers: 1) primer sequences do not contain CpGs to ensure both methylated and unmethylated alleles are amplified equally; 2) the maximum size of amplified regions is ideally less than 300bp and no more than 430bp to reduce any bias introduced from bisulfite treatment-induced DNA fragmentation; 3) amplified regions contain a minimum of 5 CpGs, and 4) primers yield a single PCR product when tested on bisulfite-treated genomic DNA (Suppl. Fig. 1A-B; Suppl. Table 2). We also designed control primers against regions consistently unmethylated (promoter and 5’end of *Sox2* and *Klf4* genes) and methylated (intronic CpG-rich region of the *Pcdha* gene cluster and last exon of the *Prickle1* gene) in mouse ESCs to control for bisulfite conversion efficiency (Suppl. Table 2). All primers also included a random 8 nucleotide barcode to enable post-sequencing data deduplication and adapter sequences to allow for efficient library construction. The IMPLICON method consists of two PCR reactions. The first PCR is an individual reaction for each primer pair which are then pooled together by sample, followed by a second PCR using barcoded Illumina adapters for sample identification (Fig. 1C, see Materials and Methods). Typical bioanalyzer traces after the first and second PCRs are displayed in Suppl. Fig. 1C. The final set of 19 primer pairs generate PCR products sampling a total of 245 CpGs (range from 5 to 23, on average 13 CpGs per amplicon) (Suppl. Table 2). Up to 32 samples can be easily processed simultaneously to generate an amplicon library in just 2-3 days, which is subsequently sequenced on an Illumina MiSeq platform (Fig. 1C).

**Fig. 1.**
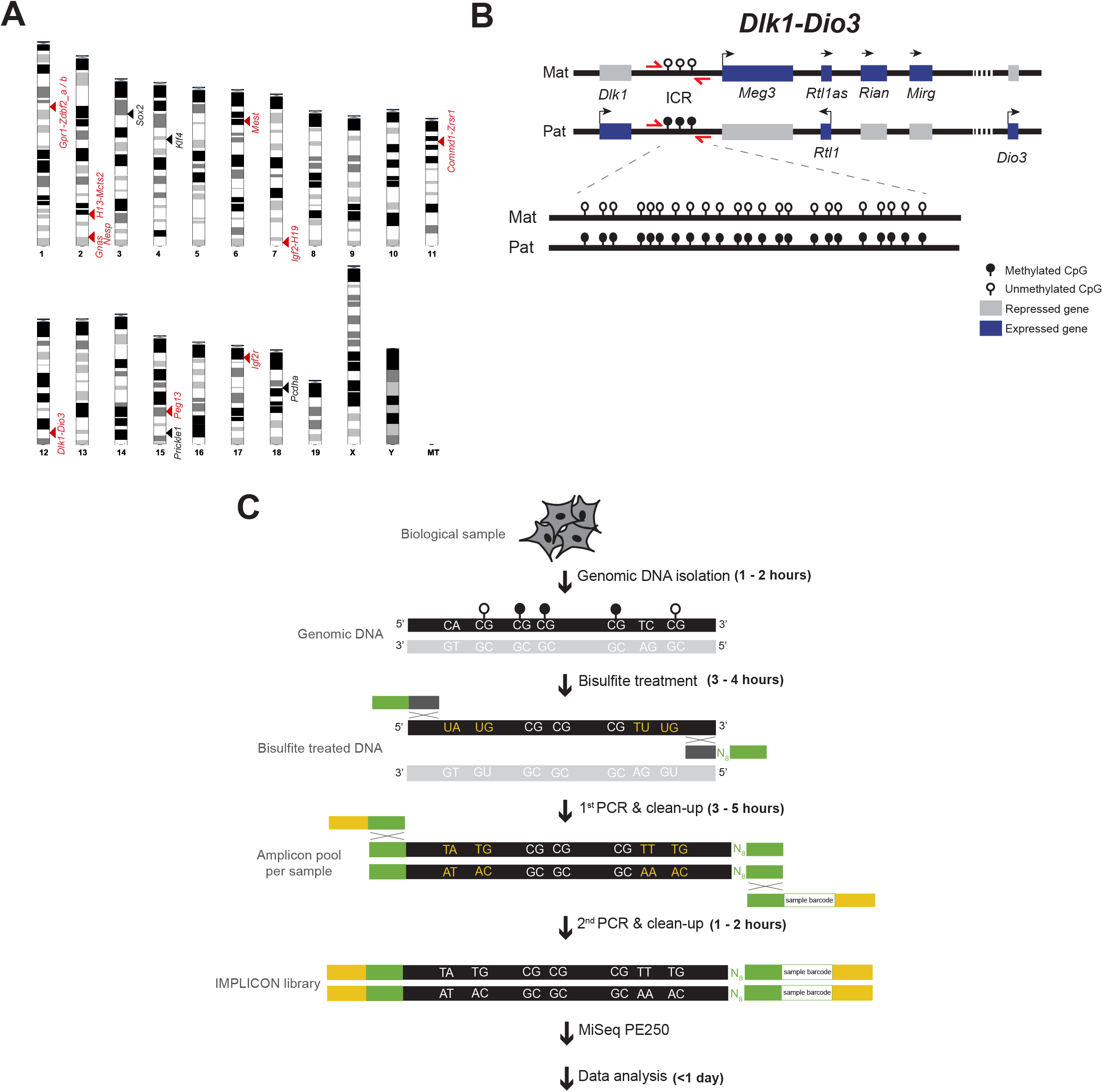
The IMPLICON method. A. Schematic view of the murine karyotype depicting the location of the regions detected by IMPLICON; black arrowheads – control regions; red arrowheads – imprinted regions. B. Schematic representation of the mouse *Dlk1-Dio3* imprinted cluster; Mat – Maternal inherited chromosome; Pat – Paternal inherited chromosome; ICR – imprinting control region; red arrows – primers to amplify *Dlk1-Dio3* ICR; genomic region is not drawn to scale. C. Brief scheme of the IMPLICON method and its approximate timeline; bisulfite conversion of genomic DNA converts unmethylated cytosines to uracils (letters in yellow), whilst methylated cytosines are retained as cytosines (in white); two rounds of PCR are then performed: the 1^st^ PCR amplifies each region for each sample separately and adds 8 random nucleotides (N_8_) for data deduplication and adapter sequences; after pooling amplicons for each biological sample, a 2^nd^ PCR completes a sequence-ready library with sample-barcodes for multiplexing; white lollipops – unmethylated CpGs; black lollipops – methylated CpGs; black DNA strand – targeted strand for amplification; light grey DNA strand – strand not targeted for amplification; Dark grey and green boxes - primers annealing the targeted strand (dark grey), containing adapter sequences (green); green and yellow - primers annealing the adapters (green), containing barcoded Illumina adapters and Illumina PE1.0 primer sequence (yellow).

### IMPLICON in inbred mice

As a proof of principle, we analysed DNA methylation levels in mouse organs (heart, liver and lung) from 3 independent adult male C57BL/6J mice of different ages (3 months, 6 months and 15 months) for which imprints are well known to be maintained (50%:50% methylated/unmethylated ratio) (23,39,40). With just one MiSeq run, we were able to examine each CpG within the selected genomic regions with an average of ~4900-fold coverage, attesting to the ultra-deep coverage of our datasets (Suppl. Table 3).

As predicted, both unmethylated (*Sox2* and *Klf4*) and methylated controls (*Pcdha* and *Prickle1*) showed, respectively, low (<~10%) or high (>~90%) levels of DNA methylation for all tested tissues in the three individuals (Fig. 2A; Suppl. Table 3). In contrast, we observed DNA methylation levels of approximately 50% for *Dlk1-Dio3* and *Gnas* imprinted regions that did not change as a function of the organ or age (Fig. 2A). Examining DNA methylation consistency for each read confirmed that the 50% methylation levels reflected an equal mix of unmethylated (<~10%) and methylated (>~90%) reads as expected for imprinted regions (Fig. 2B). DNA methylation levels of approximately 50% were also seen for all other imprinted regions analysed by IMPLICON (Fig. 2C, Suppl. Table 3).

**Fig. 2.**
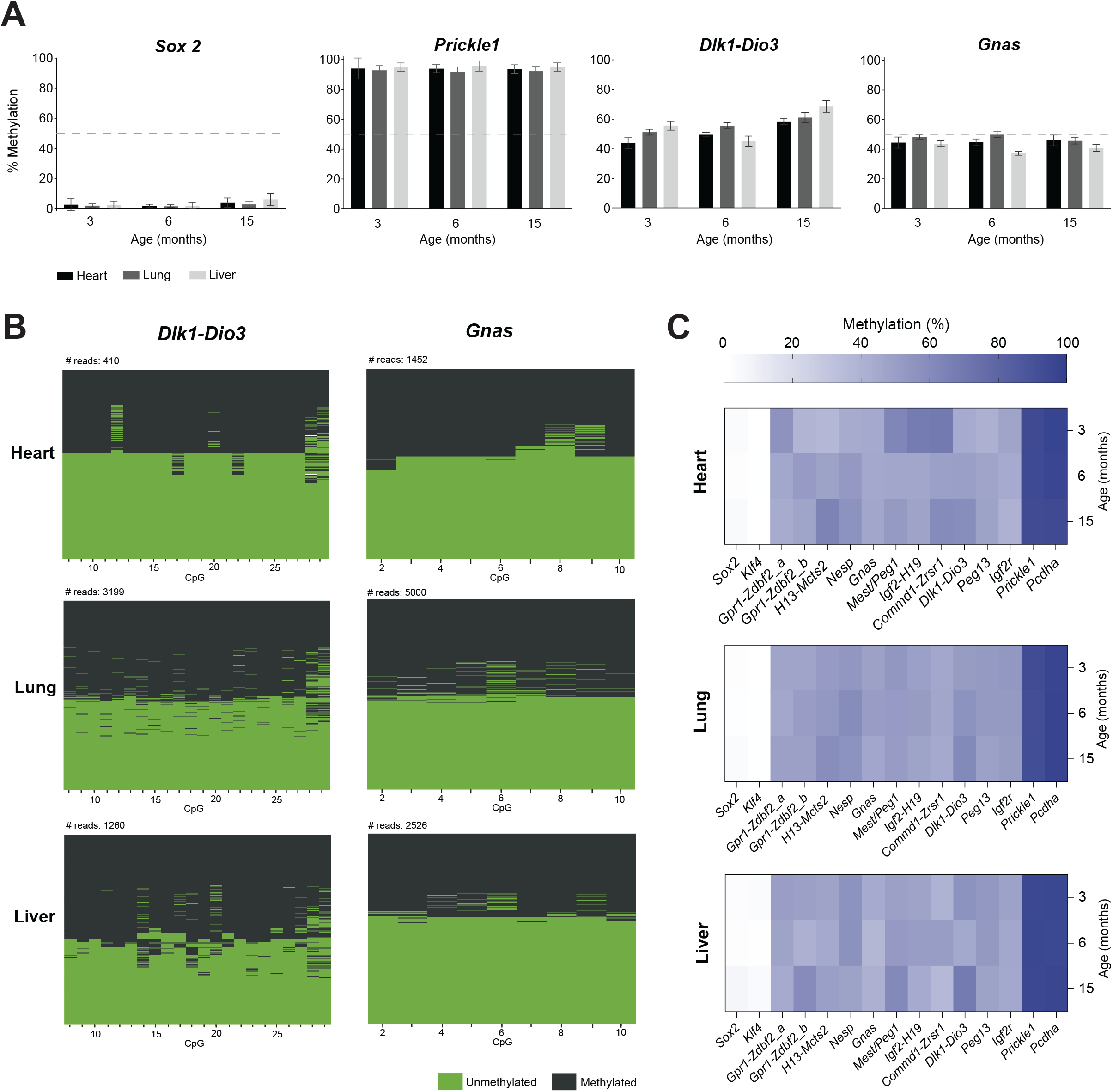
IMPLICON in adult tissues from C57BL/6J mice. A. Methylation analysis of *Sox2* (unmethylated control), *Prickle1* (methylated control) and the *Dlk1-Dio3* and *Gnas* ICRs in heart, lung and liver tissues from adult mice of different ages: 3 months, 6 months and 15 months; Graph represents the mean ± SD methylation levels measured at each CpG within each genomic region for each individual mouse; dashed line marks 50% level of methylation. B. Plot displaying methylated and unmethylated CpGs for each CpG position (in columns) in all the individual reads (in rows) for both the *Dlk1-Dio3* and *Gnas* imprinted loci in the heart, lung and liver of the 3-month-old mouse. C. Heatmap displaying average methylation levels at individual imprinted regions and controls (in columns) from adult mice of different ages (in rows) in heart, lung and liver.

### IMPLICON in hybrid mice from reciprocal crosses

To validate the parent-of-origin methylation differences, the single nucleotide polymorphisms (SNPs) of hybrid mouse crosses were used to differentiate between maternal and paternal reads. We generated reciprocal crosses of two distinct mouse strains, C57BL/6J (BL6) and CAST/EiJ (CAST), which are widely used for allele-specific studies owing to the frequent presence of SNPs (23,41) (Fig. 3A). From the original set of primers used in inbred mice, only 6 contained appropriate SNPs within the amplified region. Thus, we redesigned 10 more primer pairs according to the rules above to include SNPs not masked by bisulfite conversion (C/T SNPs were excluded) within the region of interest (see Materials and Methods; Suppl. Fig. 2; Suppl. Table 2).

**Fig. 3.**
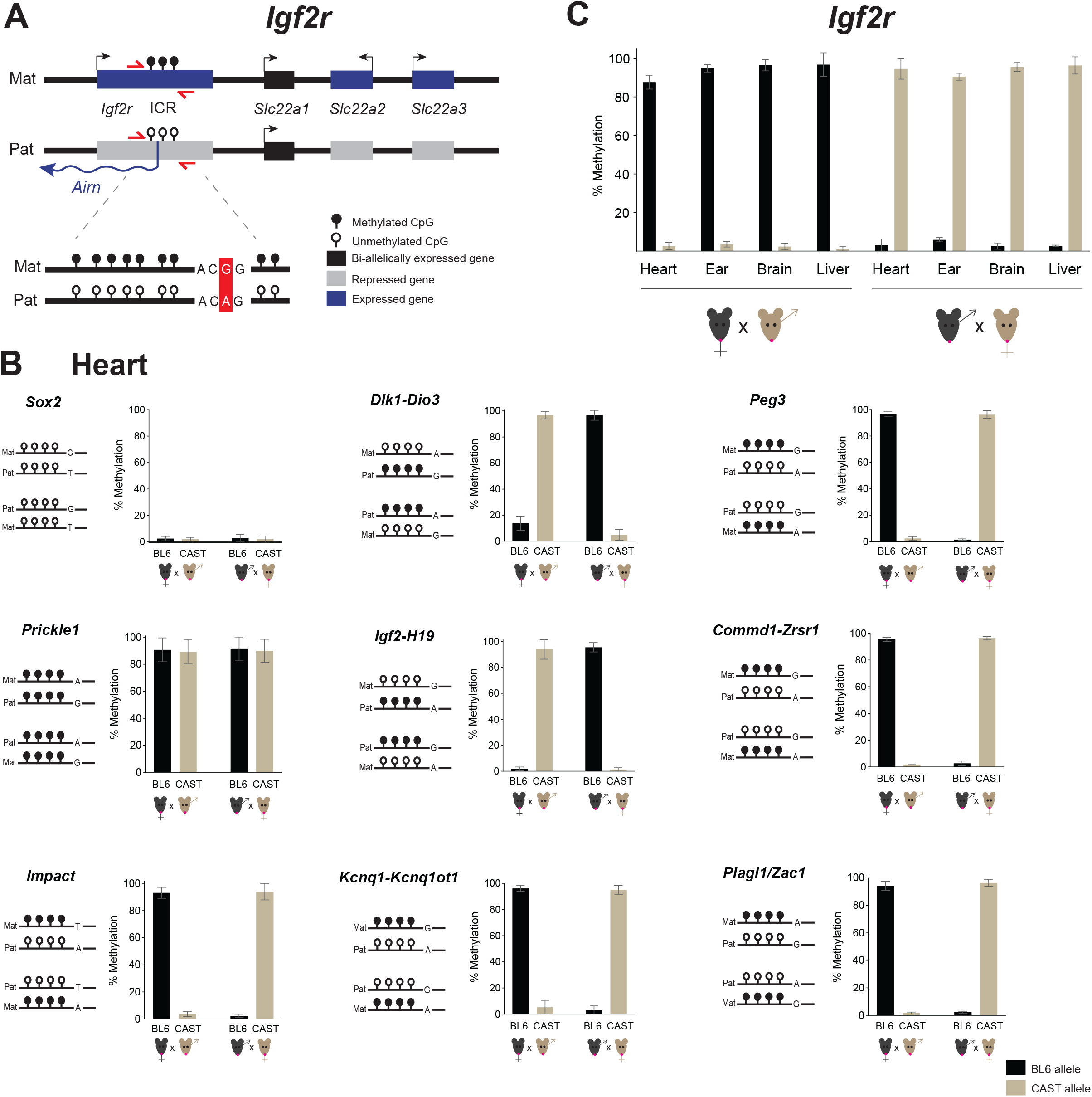
Allele-specific IMPLICON in F1 adult tissues from C57BL/6J x Cast/EiJ reciprocal crosses. A. Schematic representation of the *Igf2r* imprinted cluster; Mat – Maternally inherited chromosome; Pat – Paternally inherited chromosome; ICR – imprinting control region; red arrows – primers to amplify *Igf2r* ICR; in the scheme below, a single nucleotide polymorphism is highlighted in red; genomic region was not drawn to scale. B. Methylation analysis of *Sox2* (unmethylated control), *Prickle1* (methylated control) and ICRs of imprinted regions in the heart from F1 hybrid adult mice derived from C57BL/6J x CAST/EiJ reciprocal crosses; Graph represents the mean ± SD methylation levels measured at each CpG within different genomic regions per parental allele in the two F1 hybrid mice; Scheme on the left of each graph represents the expected methylation status of each region (white lollipops – unmethylated CpGs; black lollipops – methylated CpGs; Mat – maternal allele; Pat – paternal allele; black mice (BL6) – C56BL/6J strain; brown mice (CAST) – CAST/EiJ strain; regions are not drawn to scale. C. Methylation analysis for the *Igf2r* imprinted cluster in heart, ear, brain and liver of F1 hybrid mice from reciprocal crosses; Graph represents the mean ± SD methylation levels measured for all the CpGs within the *Igf2r* ICR in each parental allele per mouse.

We tested this allele-specific version of IMPLICON in different organs (heart, liver, brain and ear) from two F1 hybrid BL6/CAST adult male mice from reciprocal crosses (BL6 female x CAST male and vice-versa). We generated IMPLICON libraries with an average allelic coverage across the sampled CpGs that reached as high as 20,000-fold. Importantly, roughly the same proportion of reads were assigned to both the BL6 (51.26%) and CAST (48.54%) genomes, arguing against amplification bias with only 0.20% of reads left unassigned (Suppl. Table 3).

For the unmethylated (*Sox2* and *Klf4*) and methylated (*Prickle1*) controls, our results show, respectively, low (<~10%) or high (>~90%) methylation levels in both maternal and paternal hybrid alleles in the heart, but also the other tissues (Fig. 3B; Suppl. Table 3). Confident in the control regions, we turned our attention to ICRs. As exemplified in Fig. 3B, at the *Dlk1-Dio3* and *Igf2-H19* (paternally methylated) and *Peg3, Commd1-Zrsr1, Impact, Kcnq1-Kcnq1ot1* and *Plagl1/Zac1* (maternally methylated) ICRs in the heart the maternal allele was always unmethylated or methylated, respectively, independent of the strain-specific SNP. These parental allele-specific methylation patterns were unequivocally shown for all imprinted regions analysed in the four tissue samples of the same mouse (Suppl. Table 3) as exemplified for the maternally methylated *Igf2r* locus (Fig. 3C). In summary, we were able to adapt the IMPLICON method for the screening of methylation at multiple ICRs with allelic discrimination.

### Human IMPLICON

After our success in implementing IMPLICON for mouse imprinted regions, we created a human version of IMPLICON. Published methylome data from human oocytes and sperm (35) were analysed to accurately determine the genomic coordinates of gDMRs. We designed 16 primer pairs covering 14 human imprinted clusters (12 oocyte gDMRs and 2 sperm gDMRs) applying the rules above. As controls for bisulfite conversion, we included amplicons targeting regions fully unmethylated (promoter and TSS of *KLF4*) or methylated (last exon of *RHOG*) in primed human ESCs (hESCs) (Suppl. Fig. 3; Suppl. Table 2).

We first tested these primers in blood samples from two healthy individuals. Once again, we obtained high coverage for the CpGs analysed (average of approximately 6,500-fold). Our unmethylated and methylated controls showed generally low levels of DNA methylation for *KLF4* (<~10%) and high levels of methylation for *RHOG* (>~90%) (Fig. 4A). As expected, all the gDMRs inspected showed methylation levels of around 50% (Fig. 4A) reflecting an equal mix of methylated (>~90%) and unmethylated (>~10%) reads in accordance with normal imprinting patterns (Fig. 4B). Overall, these results suggest that our IMPLICON approach is suitable to look at multiple human imprinted regions.

**Fig. 4.**
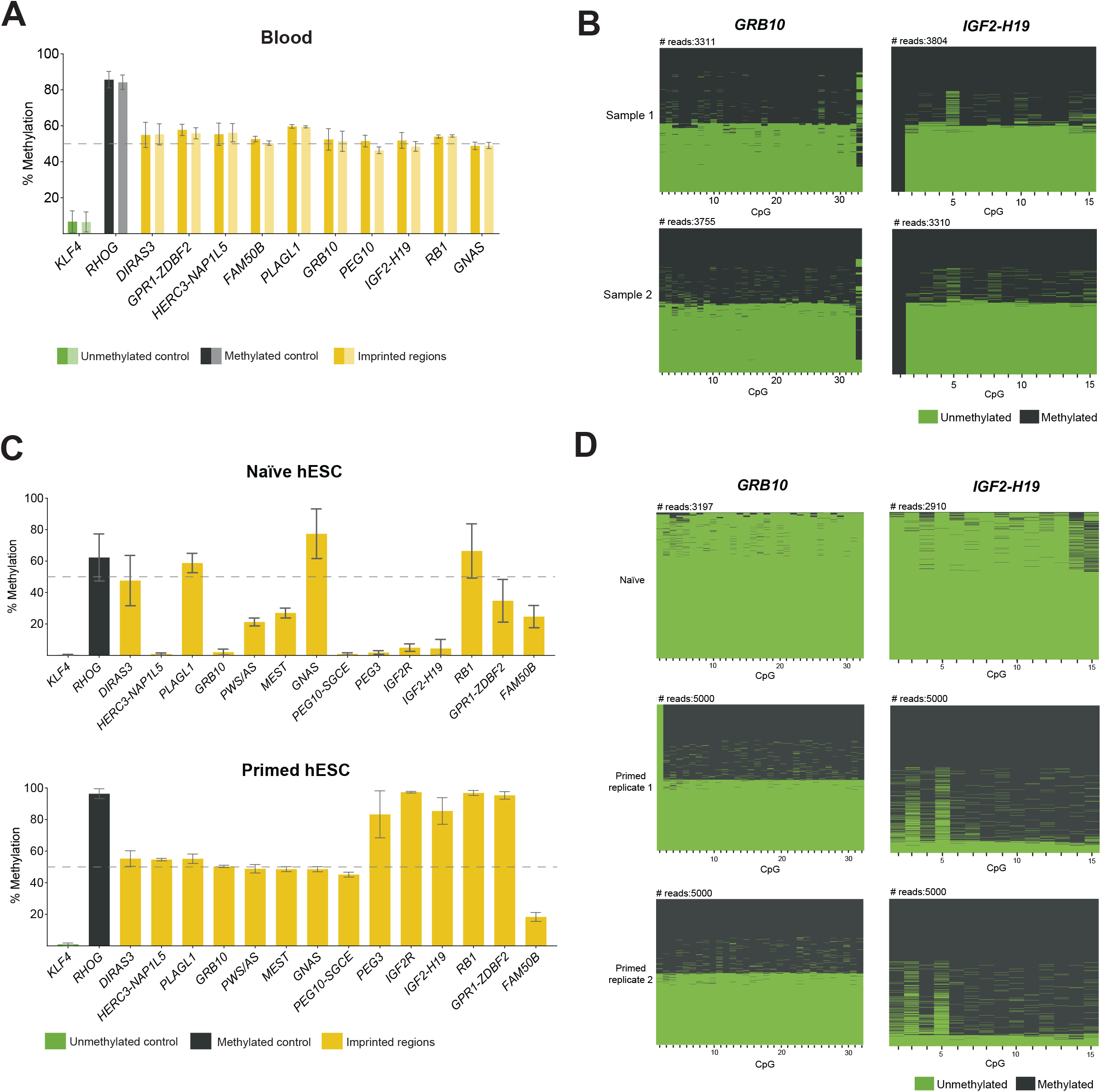
Human IMPLICON in blood and in naïve and primed hESCs. A. Methylation analysis of the *KLF4* (unmethylated control), *RHOG* (methylated control) and several gDMRs in blood samples from two independent individuals; Graph represents the mean ± SD methylation levels measured at each CpG within each genomic region for each individual blood sample (sample 1: dark colours; sample 2: light colours); dashed line marks 50% level of methylation. B. Plot displaying methylated and unmethylated CpGs for each CpG position (in columns) in all the individual reads (in rows) for *GRB10* and *IGF2-H19* for each individual blood sample (Sample 1 and 2). C. Methylation analysis of the unmethylated (*KLF4*), methylated (*RHOG*) controls and several gDMRs in naïve and primed hESCs; Graph represents the mean ± SD methylation levels measured at each CpG within each genomic region; primed hESCs shown here are an average of two replicates, whereas for naïve hESCs only one sample was analysed; dashed line marks 50% level of methylation. D. Plot displaying methylated and unmethylated CpGs for each CpG position (in columns) in all the individual reads (in rows) for *GRB10* and *IGF2-H19* for individual hESC samples (one naïve and two primed hESC replicates).

Then, we used IMPLICON to assess imprinting fidelity in different hESC culture conditions. Human ESCs cultured in naïve conditions have globally lower levels of DNA methylation (~30% compared to 70-80% in conventional or primed hESC cultures). As a result, loss of imprinting methylation is frequently observed in naïve conditions, whilst primed hESCs better maintain imprinting fidelity (42–44). In our IMPLICON results, the unmethylated control region (*KLF4)* showed <~10% methylation as anticipated (Fig. 4C). Reflecting the expected global levels of DNA methylation, the *RHOG* methylated control region showed higher (>~90%) levels of DNA methylation in primed hESCs, in comparison to naïve hESCs (~50%) (Fig. 4C). Of the 14 imprints analysed, 8 showed the expected 50% DNA methylation levels in primed hESC cultures, whereas only at 3 imprinted loci (*DIRAS3*, *PLAGL1* and *RB1*) methylation was maintained in naïve hESCs (Fig. 4C). Naïve hESCs tend to lose DNA methylation, with 10 imprinted regions having less than the 40-60% expected DNA methylation levels, whilst only *FAM50B* loses methylation in primed hESCs (Fig. 4C). This was reflected appropriately in the number of fully methylated and unmethylated reads at the *GRB10* locus: primed hESCs presented the same proportion of these reads in two biological replicates, while only fully unmethylated reads were seen for naïve hESCs (Fig. 4D). In primed conditions, 5 regions had close to 100% methylation (e.g., *IGF2-H19*), with only 1 hypermethylated region (*GNAS*) seen in naïve hESCs (Fig. 4C). *IGF2-H19* is a perfect example of a region fully methylated in primed conditions and completely unmethylated in naïve conditions (Fig. 4D). In summary, our analyses show that IMPLICON can be used successfully to identify imprinting errors in hESC cultures and furthermore highlights the importance of checking imprinting fidelity in hESC lines, including those in primed conditions.

Next, we ran our human IMPLICON on dermal fibroblasts and corresponding human induced pluripotent stem cell (hiPSC) lines previously generated from an Angelman patient and a healthy individual (33) (Fig. 5A) to search for putative imprinting defects often found in hiPSCs. As predicted, our unmethylated and methylated controls showed low (<~10 %) and high levels (>~90%) of DNA methylation, respectively (Suppl. Table 3). We then screened for the *SNURF* TSS-DMR at the PWS/AS cluster which is only methylated on the maternally inherited allele and is deleted in the Angelman patient-derived cells (Fig. 5A). While the healthy fibroblasts and hiPSCs showed the expected ~50% methylation levels, the Angelman-derived cells presented ~0% methylation, consistent with the absence of the methylated maternal *SNURF* TSS-DMR region (Fig. 5A-B). For the other imprinted regions sampled, we found values around the expected ~50% methylation in healthy and Angelman fibroblasts (Fig. 5C; Suppl. Table 3). A remarkable exception was the *IGF2R* int2-DMR, the gDMR at the *IGF2R* imprinted locus, presenting ~50% methylation in Angelman patient fibroblasts (and iPSCs), but >~90% methylation in the healthy fibroblasts (and correspondent iPSCs). Interestingly, imprinting at this region is known to be polymorphic and to differ from individual to individual (45).

**Fig. 5.**
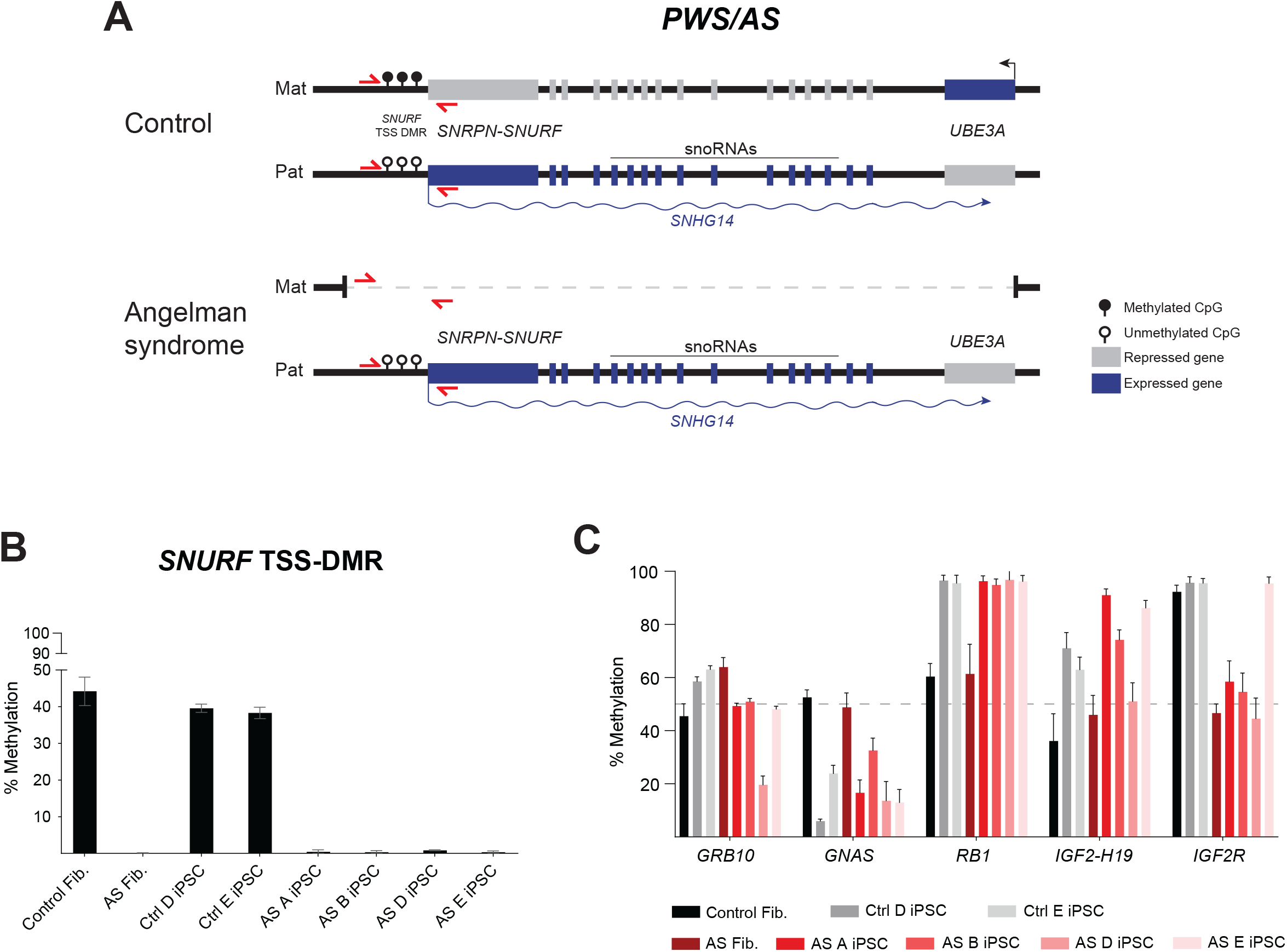
Human IMPLICON in donor fibroblasts and hiPSCs from an Angelman patient and healthy donor. A. Schematic representation of the PWS/AS locus on chr15q11-13 in human neurons, in control (healthy donor) and Angelman patient (harbouring a 4.8 Mb deletion of the maternal PWS/AS locus); *SNURF* TSS-DMR is the ICR of this region; red arrows – primers to amplify *SNURF* TSS-DMR; genomic region was not drawn to scale. B. Methylation analysis of the *SNURF* TSS-DMR in donor fibroblasts and hiPSCs from an Angelman patient and healthy donor; Graph represents the mean ± SD methylation levels measured at each CpG within the *SNURF* TSS-DMR for each sample. C. Methylation analysis of several imprinted regions (*GRB10*, *GNAS*, *RB1*, *IGF2-H19* and *IGF2R*) in donor fibroblasts and hiPSCs from an Angelman patient and healthy donor; Graph represents the mean ± SD methylation levels measured at each CpG within the *SNURF* TSS-DMR for each sample.

In contrast to stable imprinting associated with somatic cells, we observed many methylation aberrations at gDMRs for many of the imprinted clusters in hiPSCs (Fig. 5C; Suppl. Table 3). In addition to hESCs, imprinted defects have been broadly associated with hiPSCs (46–48) and are one of the major concerns for the downstream applications of these cells (49,50). A few of the imprinted loci showed no or minor abnormalities (e.g. *DIRAS3*, *GPR1-ZDBF2, MEST*, *PEG10-SGCE, GRB10*), while others show consistent hypomethylation (e.g., *GNAS*) or hypermethylation (e.g., *RB1* and *PEG3*). Furthermore, several loci displayed considerable variation from iPSC-to-iPSC line (e.g., *NAP1L5*, *FAM50B*, *PLAGL1* and *IGF2-H19*) (Fig. 4C; Suppl. Table 3). In comparison to previous reports on imprinted defects in hiPSCs that used methylome techniques (47,48), we obtained overlapping results showing a consistent tendency for hypomethylation (e.g. *PLAGL1* and *GNAS*), hypermethylation (e.g. *PEG3*), as well as, normal maintenance of imprinting (e.g. *MEST* and *PEG10*) (Suppl. Table 4). The *HERC3-NAP1L5* locus shows divergent results, however, no consistent results were observed for this locus among the previous reports (Suppl. Table 4). Overall, our IMPLICON technique identified methylation defects in gDMRs of hiPSCs consistent with previous reports highlighting its potential application in identifying imprinted defects in the context of human imprinted regions.

## DISCUSSION

We developed the new method IMPLICON to examine DNA methylation patterns at imprinted regions with an unprecedented coverage. We designed a set of primers to study murine imprinted clusters with allelic resolution, as well as human imprinted clusters. This method surpasses many shortcomings, such as time and cost of other methodologies to look at parental allele-specific methylation (Suppl. Table 1). We believe IMPLICON will provide an added value to the imprinting community and could become the gold standard for methylation inspection at multiple imprinted regions for both research and diagnostics.

Since IMPLICON is an NGS method assessing multiple imprinted regions and handling several samples in a single MiSeq run, it outperforms the traditional bisulfite sequencing and pyrosequencing methods commonly used to look at methylation in imprinted regions, being much less laborious and more high-throughput. IMPLICON yields considerably more complete datasets with over 100-fold increment in the number of amplicons analysed (compared to bisulfite sequencing) over a longer stretch of CpGs (compared to pyrosequencing). IMPLICON also has several advantages for methylation inspection at ICRs when compared to any current methylome methods (Suppl. Table 1). First, the costs associated are strongly reduced with enough reads at imprinted regions being guaranteed for multiple samples with a relatively modest sequencing depth. Second, our approach results in an oversaturated genomic coverage (>1000-fold) at imprinted regions with nucleotide and allelic resolution that is not matched by any of the other methods for the same number of sequencing reads. Finally, given the reduced costs and ultra-deep genomic coverage, IMPLICON could be easily scalable to include more genomic regions of interest and more samples.

The mouse remains the favourite animal model for studying the underlying mechanisms of imprinting regulation (reviewed in (1)). For example, the use of mouse models allowed the identification of ZFP57 and ZFP445 KRAB zinc-finger proteins as fundamental protection factors of methylation imprints at critical developmental stages (51,52). Therefore, inspection of methylation remains highly relevant for murine imprinted regions. Our initial set of primers designed on the murine reference genome (C57BL/6) consisted of 15 primers covering 9 imprinted regions. Of note, we have previously used a subset of these primers to report loss of methylation at imprinted regions in 2C-like (MERVL^+^Zscan4^+^) mouse ESCs (53), attesting for the utility of our method. The additional primers designed for allele-specific IMPLICON are also suitable to analyse imprinted loci in C57BL/6 inbred mice, which increased the number of primers to 25 covering a total of 15 imprinted clusters. This set of IMPLICON primers could, therefore, be used to look at imprinting maintenance in particular circumstances (e.g. environmental insult or genetic ablation) in cells or animals of the C57BL/6 background. Most of the primer pairs will also be suitable for imprinting analysis of other phylogenetically close strains commonly used as laboratory mice (e.g. 129/SvJ).

For imprinting studies, the use of reciprocal crosses between genetically distant mouse strains is of particular utility, since it allows for allele-specific DNA methylation and gene expression analyses based on DNA sequence polymorphisms (23,41). Importantly, our method preserves allelic information for the most commonly used BL6 x CAST cross and, moreover, it does so with ultra-deep allelic coverage (~20,000–fold was achieved). Allele-specific IMPLICON has now been optimized for 13 ICRs and future work will surely expand this set for the rest of ICRs in this hybrid cross. This could also be envisioned for other hybrid crosses (e.g. BL6 vs *Mus musculus molossinus* JF1) (54), commonly used in imprinted studies using our defined criteria for primer design (see Materials and Methods). With the ultra-deep allelic coverage achieved, we believe our method will be better at discerning subtle parental allele-specific methylation changes as a result of environmental perturbations, pathological conditions or ageing, which might never have been sufficiently appreciated using other less powerful imprinting assays (Suppl. Table 1).

Diagnostics in human medicine is undoubtedly an area where analysis of imprinting methylation is important. Besides the 8 syndromes currently characterized for which the affected imprinted loci have been identified, some patients have recently been shown to display multi-locus imprinting disturbances (MLIDs). MLIDs are characterized by epimutations in several imprinted loci and clinical manifestations of, at least, one imprinting disorder (reviewed in (11)). Screening for MLIDs, that might remain underdiagnosed, is an obvious application for our human IMPLICON method which currently covers most of the human imprinted regions and could be easily extended to all regions in the near future. Moreover, our IMPLICON method provides an easy and quick diagnostic tool not only for MLIDs, but also for other human conditions where altered imprinting is expected to be implicated, namely foetal growth restriction or cancer.

Another instance where inspection of imprinting methylation is becoming important is in stem cell biology. Indeed, genomic imprinting has been shown to gain distorted patterns through stem cell conditions and upon reprogramming of somatic cells into hiPSCs. This creates an epigenetic obstacle for their correct use in disease modelling and their application in regenerative medicine (33,46–49). In contrast to blood samples and primary dermal fibroblast, hESCs and hiPSCs exhibit several imprinting defects consistent with the reports in the literature (Suppl. Table 4) (50). This was exacerbated when hESCs were grown in naïve conditions, where loss of methylation in imprinted regions followed the globally reduced levels of DNA methylation typical of cells grown in these culture conditions (42,44). Our results show the ability of the IMPLICON method to detect these methylation deficiencies in human stem cells. Since epigenetic stability in stem cells remains an issue and genomic imprinting provides an excellent thermostat of epigenetic fidelity, IMPLICON emerges as a simple and fast method for routine screening of human ESC/iPSCs ahead of their use in downstream applications.

In summary, we present a rapid method to measure imprinting methylation in a high-throughput and cost-effective manner that surpasses the limitations of other high-throughput sequencing methods when imprinting inspection is at stake (Suppl. Table 1). With further developments, IMPLICON could easily cover all known ICRs. Importantly, the guidelines and rules presented are extensive to screen DNA methylation profiles in any other genomic regions where high coverage is desired. With unprecedented coverage and nucleotide resolution, IMPLICON could become a gold-standard method to profile imprinting methylation in laboratory and clinical settings where aberrant imprinting has been or is believed to be implicated.

## ACKNOWLEDGEMENTS

We would like to thank members of Wolf Reik’s lab and S.T.d.R.’s team for the helpful discussions. We would like to thank Wolf Reik for useful discussions and for critical reading of the manuscript. We would like to also thank Ruslan Stroganstev for helpful insights about genomic imprinting, Tom Stubbs for the help with primer design and Babraham Institute Bioinformatics for bioinformatic advice. We thank Bethan Hussey and Elizabeth Easthope at Sanger Institute and Kristina Tabbada and Nicole Forrester at Babraham Institute for assistance with high-throughput sequencing.

