## Supplementary Information for "IMPLICON: an ultra-deep sequencing method to uncover DNA methylation at imprinted regions"

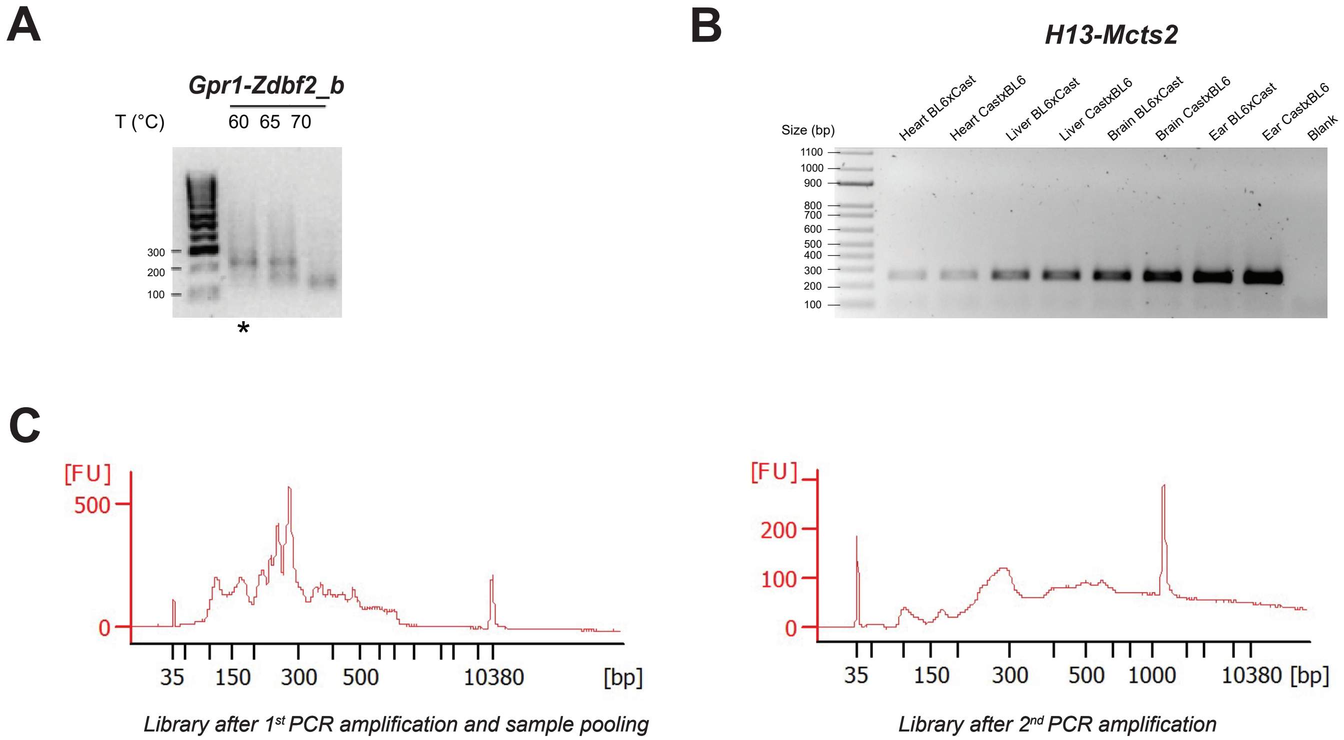


### Supplementary Figure 1

### *Mouse_allelic-specific*

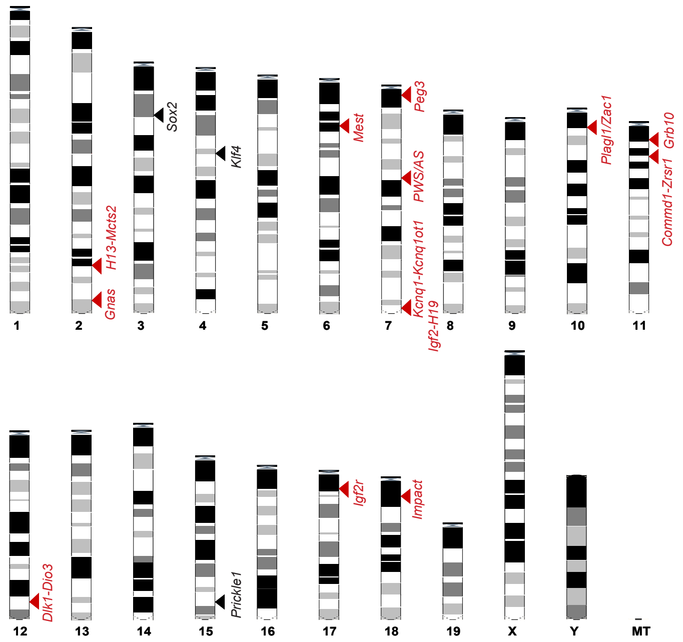


### Supplementary Figure 2

### *Human*

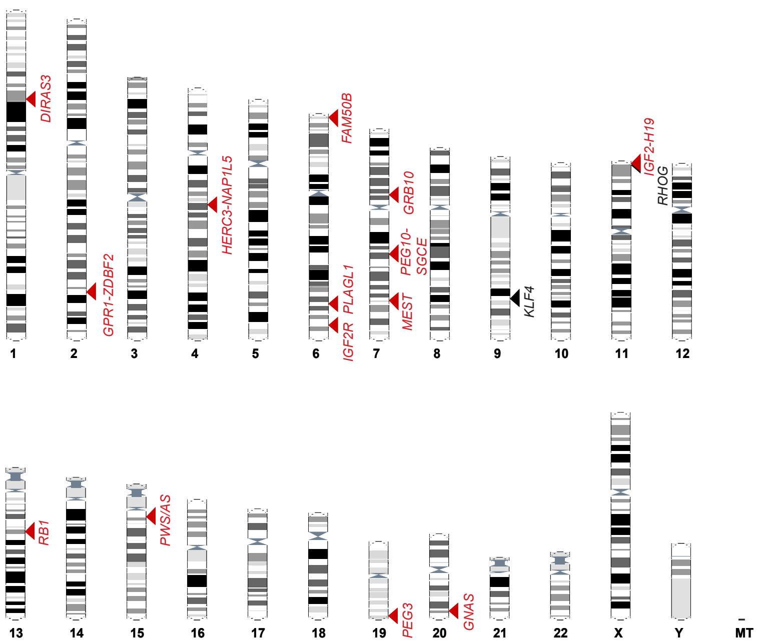


### Supplementary Figure 3

**SUPPLEMENTARY FIGURE LEGENGS**

**Suppl. Fig. 1 – Steps of quality control of the IMPLICON method.**

A. Agarose gel displaying the primer optimization step for 1st PCR for the *Gpr1_Zdbf2_b* primer pair on *Gpr1-Zdbf2* locus; primer pair was tested with different annealing temperatures in mouse ESCs; *represents conditions chosen for the IMPLICON run.
B. Agarose gel displaying an example of individual PCR reactions after the 1st PCR step, amplifying *H13-Mcts2* locus in different tissues (heart, liver, brain, ear) from F1 hybrid mice; Blank represents a negative water control.
C. Library profiles obtained upon analysis on Agilent bioanalyzer after 1st PCR amplification and sample pooling (left) and after a 2nd PCR amplification and clean-up (right).

**Suppl. Fig. 2 – Schematic view of the murine karyotype depicting the location of the regions detected by allele-specific IMPLICON; black arrowheads – control regions; red arrowheads – imprinted regions.**

**Suppl. Fig. 3 – Schematic view of the human karyotype depicting the location of the regions detected by the human version of IMPLICON; black arrowheads – control regions; red arrowheads – imprinted regions.**

| **Technique** | **Throughput** | **Costs** | **Time** | **Bisulfite conversion** | **Single CpG resolution** | **Advantages for imprinting analysis** | **Disadvantages for imprinting analysis** | **References** |
|---|---|---|---|---|---|---|---|---|
| WGBS - Whole genome bisulfite sequencing or MethylC-seq | Low | Very High | 2-4 weeks | Yes | Yes | Virtual representation of all imprinted regions at single base resolution | Low coverage of imprinted regions (~15x); Elaborate bioinformatics | Xie et al., 2012 |
| RRBS - reduced representation bisulfite sequencing | Low | High | 2-4 weeks | Yes | Yes | Greater coverage than WGBS at the imprinted regions represented at single base resolution | Absence of few imprinted clusters due to low genome coverage (10%); Elaborate bioinformatics | Stelzer et al., 2013 |
| MeDIP-seq - Methylated DNA Immunoprecipitation sequencing | Low | High | 2-4 weeks | No | No | No 5hmC detection | Low base resolution; biases towards hypermethylated regions | Proudhon et al., 2012 |
| Illumina Infinium MethylationEPIC array | Medium | Medium | 1-2 weeks | Yes | No | Catalog of MethylationEPIC probes for fast screening of human imprinted regions | Relative measurement; High signal to noise ratio | Hernandez Mora et al., 2018 |
| Long-read Nanopore Sequencing | Low | Very High | 2-4 weeks | No | Yes | Long reads; Direct 5mC detection | Low coverage of imprinted regions (~10x); Elaborate bioinformatics | Gigante et al., 2019 |
| IMPLICON | High | Low | < 1 week | Yes | Yes | Ultra-deep genomic coverage (>1000 reads) at single allele and single base pair resolution | A few imprinted regions to be added | This work |

**Suppl. Table 1 - Advantages and disadvantages of current high-throughput methods to quantify DNA methylation at imprinted regions.**

Abbreviations: 5hmC – 5’hydroxylmethyl-cytosine; 5mC – 5’methyl-cytosine.

| Imprinted cluster | Disease associated | Nazor et al., (2012): Infinium 450K BeadChip | Ma et al., (2014): Infinium 450K BeadChip | This study: IMPLICON |
|---|---|---|---|---|
| *DIRAS3* | - | Tendency for Hypermethylation | Tendency for Hypermethylation | Normal |
| *GPR1/ZDBF2* | - | - | - | Normal |
| *HERC3-NAP1L5* | - | Tendency for Hypermethylation | Normal | Tendency for Hypomethylation |
| *FAM50B* | - | - | - | Tendency for Hypomethylation |
| *PLAGL1* | Transient Neonatal Diabetes Mellitus | Tendency for Hypomethylation | Tendency for Hypomethylation | Tendency for Hypomethylation |
| *IGF2R* | - | - | - | Polymorphic imprinting |
| *GRB10* | - | Tendency for Hypomethylation | Normal | Rare Hypomethylation |
| *PEG10-SGCE* | Silver-Russell syndrome | Normal | Normal | Normal |
| *MEST* | - | - | Normal | Normal |
| *IGF2-H19* | Silver-Russell syndrome; Beckwith-Weideman syndrome | Tendency for Hypermethylation | Normal | Tendency for Hypermethylation |
| *RB1* | - | - | - | Hypermethylation |
| *PWS-AS* | Prader-Willy syndrome; Angelman syndrome | Tendency for Hypomethylation | Normal | Normal |
| *PEG3* | - | Hypermethylation | Hypermethylation | Hypermethylation |
| *GNAS* | Sporadic pseudohypoparathyreoidism Ib | Tendency for Hypomethylation | Rare Hypomethylation | Hypomethylation |

**Suppl. Table 4 - Comparative analysis of methylation defects at imprinted regions in human induced pluripotent stem cells from three methylome studies.**
